# Grey matter reshaping of language-related regions depends on tumor lateralization

**DOI:** 10.1101/2023.02.02.526219

**Authors:** Lucia Manso-Ortega, Laura De Frutos-Sagastuy, Sandra Gisbert-Muñoz, Noriko Salamon, Joe Qiao, Patricia Walshaw, Ileana Quiñones, Monika M. Połczyńska

## Abstract

A brain tumor in the left hemisphere can decrease language laterality as assessed with fMRI. However, it remains unclear whether or not this decreased language laterality is associated with a structural reshaping of the grey matter, particularly within the language network. Here, we examine if the disruption of language hubs exclusively affects macrostructural properties of contralateral homologues (as suggested by previous research), or whether it affects both hemispheres. This study uses voxel-based morphometry applied to high-resolution MR T1-weighted MPRAGE images from 31 adult patients left-dominant for language. Eighteen patients had brain tumors in the left hemisphere, and 13 had tumors in the right hemisphere. A cohort of 71 healthy individuals matched on age and sex was used as a baseline. We defined 10 ROIs per hemisphere known to subserve language function. Two separate repeated-measures ANOVAs were conducted with the volume per region as the dependent variables. For the patients, tumor lateralization (right versus left) served as a between-subject factor. The current study demonstrated that the presence of a brain tumor generates a global volumetric change affecting left language regions and their contralateral homologues. These changes are mediated by the lateralization of the lesion. Our findings suggest that compensatory functional mechanisms are supported by the rearrangement of the grey matter, although future longitudinal research should determine the temporal course of such changes.

## Introduction

The presence of a brain tumor can impair essential cognitive abilities, such as language. Consequently, the brain may reorganize to compensate for the presence of the lesion (Herbet et al. 2016). Patients with brain tumors can serve as an ideal pathological model to enhance our understanding of lesion-dependent plasticity. When language hubs are damaged in this population, functional compensation involving the recruitment of ipsilesional or contralesional regions has been observed to support recovery (Ille et al. 2019; Krieg et al. 2013; Li et al. 2019; Quiñones et al. 2021; Połczyńska et al. 2021). However, it remains poorly understood how the brain structurally responds to tumors harboring language hubs. While the scarce available evidence shows that patients with brain tumors in the language network had increased grey matter (GM) volume in contralateral regions (Almairac, Duffau, and Herbet 2018; Hu et al. 2020; Yuan et al. 2020), we do not yet know if structural changes are induced more globally, including ipsilaterally (Pasquini et al. 2022). The goal of this work is to examine how tumor laterality affects language structures and their right hemisphere homologues. To fulfill this goal, we examined GM volume within the language network in the left hemisphere and its right homologues in patients with brain tumors affecting language hubs. Further, we determined whether changes in macrostructural properties are evidenced by variations in GM that could constitute a structural compensation mechanism.

Brain tumors located in the left language-dominant hemisphere can decrease language laterality, as shown by functional magnetic resonance imaging (fMRI) assessments (Połczyńska et al. 2021; Batouli et al. 2016). Decreased functional laterality has been linked to weaker activation during language tasks in structures close to the lesion (Kristo et al. 2015), increased contralesional activity in right language homologues, or both (Partovi et al. 2012; Petrovich et al. 2004; Ulmer et al. 2004; Krieg et al. 2013; Chivukula et al. 2018). In contrast, tumors located in the right hemisphere (which is not dominant for language) have been shown to have little to no effect on language activation during fMRI tasks (Połczyńska et al. 2021; Ille et al. 2019; Wang et al. 2013). It is yet to be determined whether functional compensation is accompanied by macrostructural reshaping within the language network.

At present, plasticity in patients with brain tumors remains fairly understudied. Resectability rates of cortical and subcortical structures point to a far greater plastic potential of GM than white matter (WM) (Ius et al. 2011; Sarubbo et al. 2015, 2020). However, to our knowledge, only 3 studies have used voxel-based morphometry (VBM) with GM volume as an index of structural plasticity in patients with brain tumors. The first study applied VBM to measure GM volume in 47 patients with gliomas in the left insula and 37 individuals with gliomas affecting the right insula (Almairac, Duffau, and Herbet 2018). The authors noted that both patient groups displayed a marked increase in the volume of the contralesional insula, suggesting that the unaffected insula was recruited via an intact corpus callosum. Following a similar approach, Hu et al. (2020) investigated contralesional GM volumes associated with cognition in patients with temporal lobe glioma. In the preoperative assessment, the authors included a sample of 8 patients with a tumor in the left temporal lobe and 9 patients with a tumor in the right temporal lobe. Compared with 28 matched healthy controls, the patients with left temporal tumors showed increased GM volume in the right inferior temporal gyrus and right superior temporal pole, whereas patients with right temporal tumors exhibited increased GM volume in the left inferior temporal gyrus. It was concluded that the increases in the homologue contralesional structures in both patient groups served as a physiologic foundation for high levels of functional compensation. More recently, structural alterations in the contralesional medial temporal lobe (MTL) were studied in 68 individuals with gliomas (Yuan et al. 2020) using VBM. Patients showed decreased GM in this region when compared to 40 healthy volunteers.

A clear limitation of the research presented above is that the studies only analyzed structural alterations in homologous brain sites that were contralesional to the tumor: one focused on the insula (Almairac, Duffau, and Herbet 2018) and two on the temporal lobe (Hu et al. 2020; Yuan et al. 2020). These regions have been implicated in several cognitive processes rather than being specific for language (Hickok and Poeppel 2007; Binder 2017). Yet, none of the studies accounted for ipsilateral regions in the left hemisphere, either those adjacent to the tumor or ipsilateral regions more distant from the lesion.

The current work examines the macrostructural properties of plasticity within the language network in patients with tumors affecting the left language-dominant hemisphere. For this purpose, GM volume was analyzed as an index of structural plasticity. We measured GM volume of previously identified language regions and their right homologues (Hickok and Poeppel 2004, 2007; Binder 2017). We hypothesized that patients with tumors in the left-language dominant hemisphere would show macrostructural changes in all regions within the language network, including the right homotopic regions of interest and the close and far left ipsilateral regions. Changes in GM volume could then be considered as a sign of structural compensation for sustaining language ability after a brain lesion. We included 2 control groups: (1) patients with tumors in the right non-dominant hemisphere, in whom we predicted no structural alterations (Ille et al. 2019; Połczyńska et al. 2021), and (2) 71 healthy volunteers matched on age and sex.

## Methods

### Participants

The clinical sample consisted of 31 patients (19 females, mean age = 47.6 years, SD = 13.9 years) who were diagnosed with brain tumors. The data from all patients was obtained before a planned surgery, regardless of the existence of previous surgeries for some of the patients (as can be seen in Table 1). Patients were divided into 2 groups: (1) left tumor group with 18 individuals (target group), and (2) right tumor group with 13 individuals (control group). A summary of the patient demographics and clinical characteristics can be found in Table 1. All patients were clinically classified as having left-language dominance, based on clinical conclusions from neurocognitive assessments, clinical language fMRI, and (in several cases) direct cortical stimulation. The recording of the pathological data was overseen by the Institutional Review Board at the University of California (code: K01DC016904), Los Angeles following the Code of Ethics of the World Medical Association (Declaration of Helsinki) for experiments involving humans. As we accessed and retrospectively analyzed presurgical structural MRI data from a larger published sample of patients with brain tumors, we followed the inclusion criteria described in Połczyńska et al. 2021. We included English monolingual patients who did not have aphasia significant enough to fail the initial screening examination for assessing language.

**Table 1.** Patient characteristics.

|  | **Left**<br>**tumors**<br>**(n=18)** | **Right**<br>**tumors**<br>**(n=13)** |
| --- | --- | --- |
| **Handedness** |  |  |
| Right | 89% | 38% |
| Left | 6% | 46% |
| 71)7171Ambidextrous | 6% | 15% |
| **Tumor type WHO** |  |  |
| Diffuse | 83% | 77% |
| Metastatic tumors | 11% | 0 |
| No data | 6% | 23% |
| **Previous surgery** |  |  |
| Yes | 61% | 38% |
| No | 28% | 53% |
| No data | 11% | 8% |
| **Aphasia** |  |  |
| None | 22% | 46% |
| Mild | 39% | 46% |
| Moderate | 22% | 8% |
| Severe | 6% | 0 |
| No data | 11% | 0 |

Tumor type diffuse refers to both astrocytic and oligodendrial tumors.

In addition, healthy-control data from 71 participants matched on age and sex (43 women, mean age = 44.49; SD = 12.5) was used as a reference point. Specifically, we characterized the structural relationships among regions in the typical language network to interpret potential divergent patterns of structural reshaping in the patient groups. The study protocol was conducted in accordance with the Declaration of Helsinki for experiments involving humans and was approved by the Ethics Board of the Euskadi Committee (code: 270220SM) and the Ethics and Scientific Committee of the Basque Center on Cognition, Brain and Language (BCBL). Informed consent was obtained from all participants involved in the study before the experiment.

### MRI data acquisition and lesion reconstruction

To minimize the possible bias caused by using images from different centers, the main acquisition parameters (e.g., magnet strength, model of scanner, and version of pulse sequence) were standardized. High-resolution MR T1-weighted MPRAGE images were obtained from the patients and controls using a 3T Siemens scanner (for clinical data: MAGNETOM Allegra with a 20-channel head coil; for healthy controls: a MAGNETOM Prisma with a 64-channel head coil) with 176 slices of 1-mm isotropic resolution and matrix size 256 × 256 (for clinical data: TR 1900 msec, TE 2200 msec and flip angle 9°; for healthy controls: TR 2530 msec, TE 2360 msec, and flip angle 120°). Two trained technicians manually drew the lesions slice-by-slice using the MRIcroGL (Rorden and Brett 2000) free and open-source software. They took tumor masks drawn by trained clinicians and researchers at the University of California, Los Angeles as a reference. To determine GM and WM in regions affected by the lesion and to estimate tumor volume (cm^3^) per participant, in-house MATLAB (2014b release, Mathworks, Inc, Natick, MA) codes were developed using functions from Statistical Parametric Mapping (SPM12, Welcome Department of Cognitive Neurology, London, UK) and the related toolbox. Lesion overlap maps are displayed in Figure 1.

**Figure 1.**
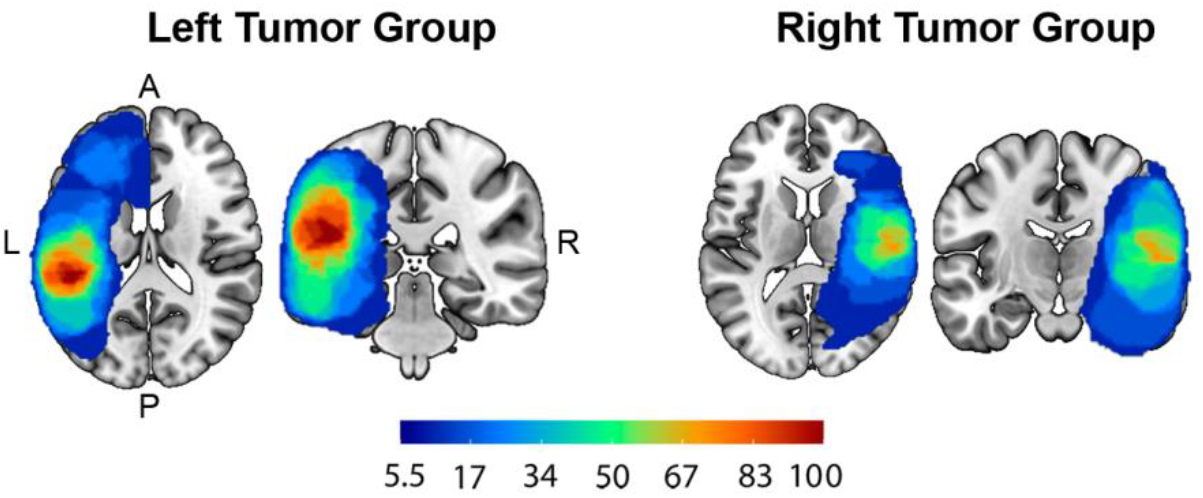
Lesion overlap per tumor group. The heatmaps – from red to blue – represent the number of overlapping tumors within the left and the right tumor groups in percentages.

### MRI analyses

For all participants, voxel-based morphometry (VBM) was performed using SPM12 in MATLAB. This semi-automatized neuroimaging method has been successfully used to quantify macrostructural brain changes – in volume or density – in longitudinal and cross-sectional studies on development and disease (Ashburner and Friston 2001). Previous results demonstrated that this type of multi-center study is methodologically feasible and reliable for the assessment of local changes in tissue integrity induced by a given pathological condition (Biswal et al. 2010; Bordin et al. 2021). The processing pipeline was standardized for patients and healthy controls as described by Ashburner and Friston (2001).

Due to differences arising from noise in the images, we removed the skull and non-brain structures for patients as a first step. Skull stripping was performed on the T1 MPRAGE images using the brain extraction function in SPM12. For the healthy participants, a manual trimming of the images was performed. Then, the images were manually reoriented and shifted to set the anterior commissure, a bundle of white matter fibers that connect the anterior lobes of the brain, as the origin. Next, T1 MPRAGE-weighted images were segmented into the GM, WM and cerebrospinal fluid (CSF) following the segmentation module in SPM12. The volumes of the native segmentations (GM, WM, and CSF) were computed and used to calculate the total intracranial volume (TIV) of each participant. To calculate the GM volume per region, we used the automated anatomical labeling (AAL) atlas (Tzourio-Mazoyer et al. 2004). The volume of each of the 116 ROIs of the AAL atlas was calculated and divided by the TIV. The use of proportionally scaled scores (volume per ROI divided by TIV) instead of using GM segmentations, minimizes potential bias due to variables that we were not able to control for and could potentially affect each individual differentially.

To investigate the impact of brain tumors on neuroplastic structural mechanisms affecting ipsilateral and contralateral language areas, 10 regions that are critically involved in language processing were selected: pars orbitalis, pars opercularis, and pars triangularis within the inferior frontal gyrus (IFG), middle frontal gyrus, middle temporal gyrus, middle temporal pole, superior temporal gyrus, superior temporal pole, supramarginal gyrus and angular gyrus (Figure 2).

**Figure 2.**
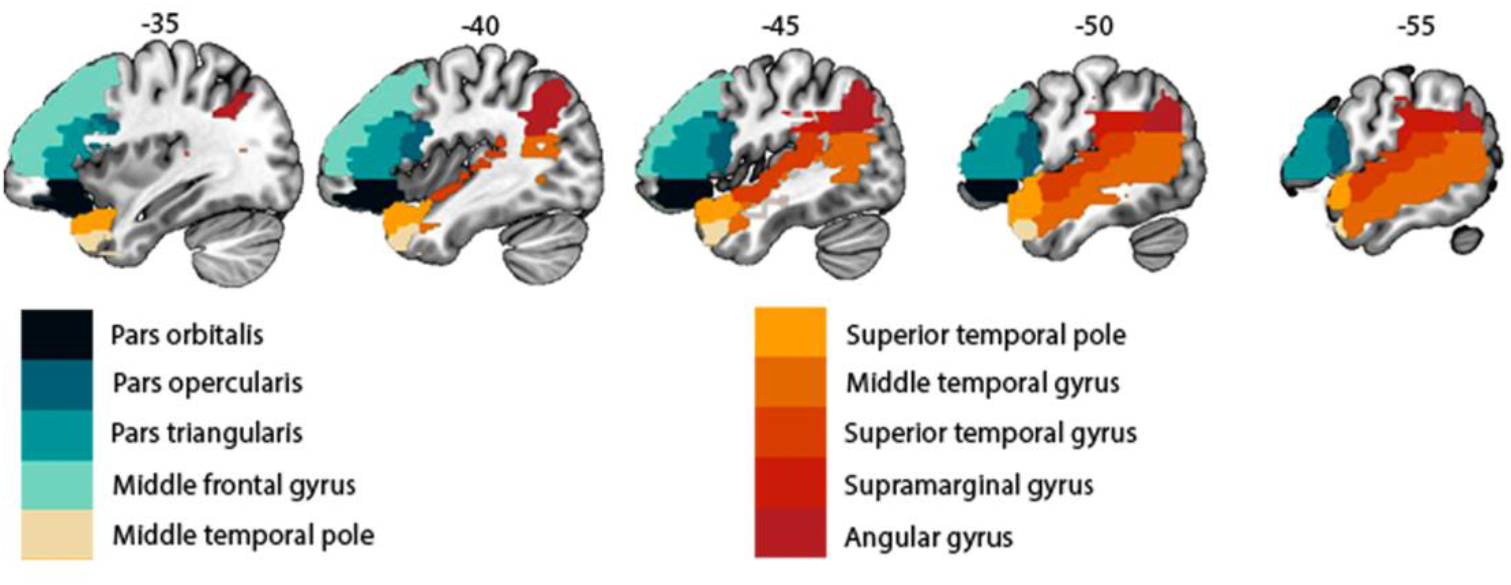
Picture created following the AAL atlas parcellation with the selected language regions known to subserve language production (i.e., pars orbitalis, pars opercularis, pars triangularis, middle frontal gyrus) and comprehension (middle temporal gyrus, middle part of the temporal pole, superior temporal gyrus, superior temporal lobe, supramarginal gyrus, angular gyrus).

### Statistical approach

Statistical comparisons were performed keeping patients and healthy controls in separate designs in order to avoid potential effects due to the differences between the scanners inherent to the different populations. A repeated measures ANOVA was performed with GM volume for the 10 ROIs weighed by the TIV (for patients: mean TIV = 1536.08; SD = 237.36; for healthy controls: mean TIV = 1419.92; SD = 158.22) as dependent variables Two within-subject factors were included: (1) ROI lateralization (right/left) and (2) Regions of interest (ROIs): 10 ROIs were considered including frontal, temporal and parietal areas. As patients had lesions affecting either the left or the right hemisphere, lesion lateralization was considered as a between-subjects factor controlling for lesion volume and tumor grade. We also included for both groups – patients and healthy controls – the age and sex as nuisance covariates. Pairwise comparisons were calculated as a post-hoc analysis applying Bonferroni correction for multiple comparisons.

## Results

### Structural reshaping in patients with brain tumors

After performing Greenhouse-Geisser sphericity correction, we found double and triple interactions (illustrated in Figure 3). We found that patients with tumors in the left hemisphere had more volume in the right ROIs (contralesional) relative to the same ROIs in the left hemisphere (ipsilesional), as shown by the double interaction between ROI lateralization (right/left) x Lesion lateralization (right/left) (F(1)=25.93, p<0.001). The triple interaction of ROI lateralization x ROIs x Lesion lateralization (F (9) =6.29, p<0.001) was also significant, so pairwise comparisons were calculated as a post-hoc analysis (Table 2). For patients in the left tumor group, after applying Bonferroni correction, 5 out of 10 structures exhibited larger volumes in the right hemisphere relative to the left: the angular gyrus, superior temporal gyrus, middle temporal pole, and the pars opercularis within the IFG. One ROI displayed a reversed result with more volume on the left than right side (pars triangularis within the IFG) (Figure 3B). Individuals with tumors in the right hemisphere had similar volume values for the left and right ROIs, with 3 larger structures on the left (contralesional) (see Figure 3C). These areas were the middle temporal gyrus, pars orbitalis, and pars triangularis. The angular gyrus was the only structure that had more volume in the right hemisphere compared to the left.

**Table 2.** Paired samples t-test for each group (R>L). Effect size is given by Cohen’s d.

| Regions | Left Tumor Group | Right Tumor Group | Healthy Controls |
| --- | --- | --- | --- |

|  | **T** | **p** | **Cohen's d** | **T** | **p** | **Cohen's d** | **T** | **p** | **Cohen's d** |
| --- | --- | --- | --- | --- | --- | --- | --- | --- | --- |
| Pars orbitalis | -1.11 | 0.282 | -0.26 | **-4.50** | <0.001* | -1.25 | **3.50** | <0.001* | 0.42 |
| Pars opercularis | **6.29** | < .001* | 1.48 | 0.55 | 0.590 | 0.15 | **55.24** | <0.001* | 6.56 |
| Pars triangularis | **-4.33** | 0.001* | -1.02 | **-3.65** | < .003* | -1.01 | **-36.11** | <0.001* | -4.29 |
| Middle frontal gyrus | 1.55 | 0.139 | 0.37 | 0.22 | 0.827 | 0.06 | **21.84** | <0.001* | 2.59 |
| Middle temporal pole | **4.13** | < .001* | 0.97 | 0.93 | 0.369 | 0.26 | **77.36** | <0.001* | 9.18 |
| Superior temporal pole | 1.88 | 0.078 | 0.44 | -1.46 | 0.171 | -0.40 | **10.45** | <0.001* | 1.25 |
| Middle temporal gyrus | 1.13 | 0.275 | 0.27 | **-4.67** | < .001* | -1.30 | **-40.38** | <0.001* | -4.79 |
| Superior temporal gyrus | **5.40** | < .001* | 1.27 | -1.66 | 0.123 | -0.46 | **76.76** | <0.001* | 9.11 |
| Supramarginal gyrus | **6.51** | < .001* | 1.53 | -0.25 | 0.811 | 0.07 | **90.24** | <0.001* | 10.71 |
| Angular gyrus | **4.85** | < .001* | 1.14 | **4.89** | <0.001* | 1.36 | **92.25** | <0.001* | 10.95 |

T values in bold represent significant comparisons after Bonferroni correction.

**Figure 3.**
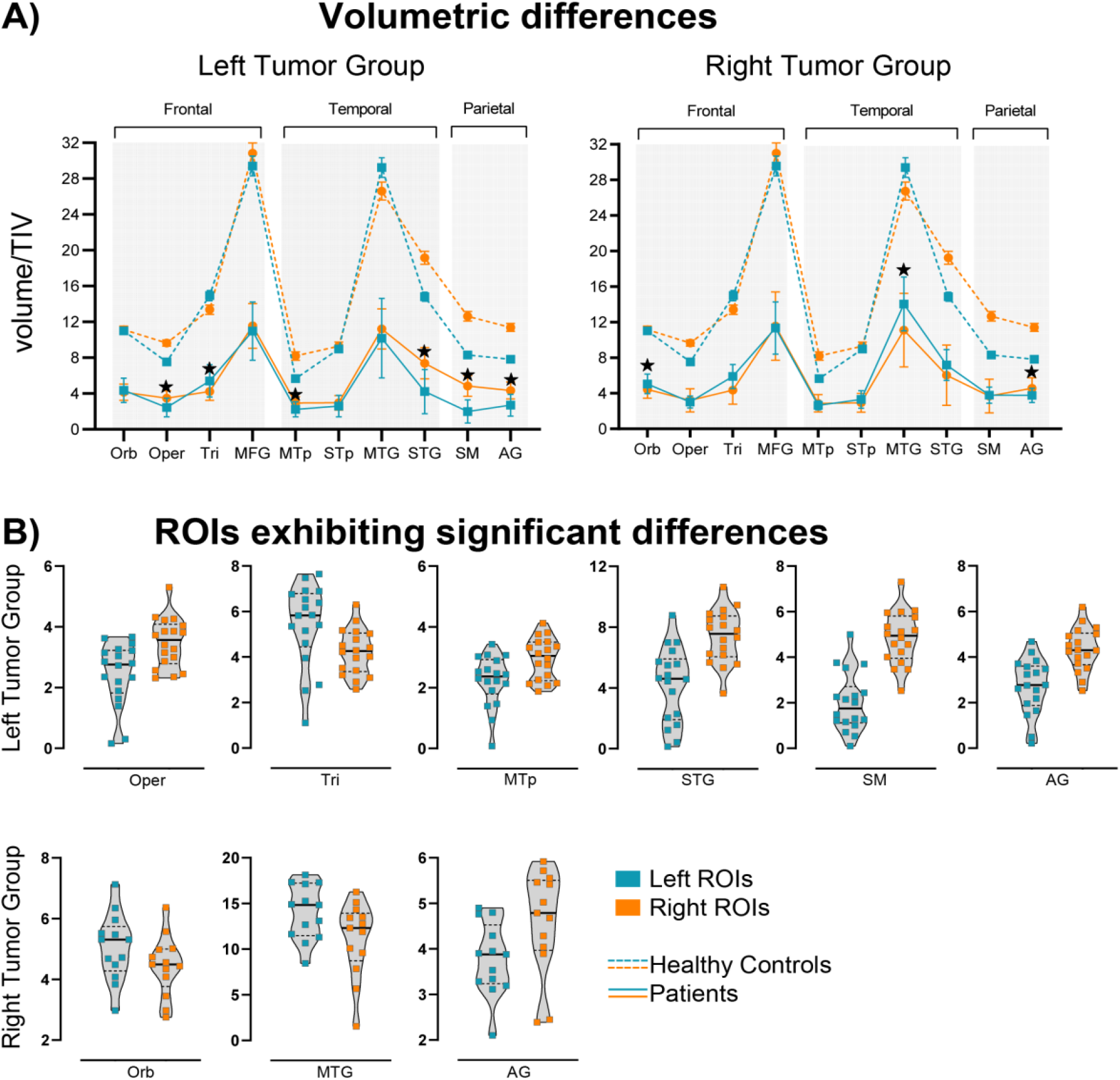
Line charts show volumetric differences for each group of patients considering the 10 ROIs (right vs. left). The ratio between volume per region and TIV values are also shown for healthy controls as indicated by dotted lines. Stars indicate that the comparison between contralateral regions reached significance after Bonferroni correction (p<0.002).

### Structural lateralization of the language network in healthy participants

We report a main effect of ROI lateralization (F (1) = 139.025, p<0.001) and, after Greenhouse-Geisser sphericity correction, a double interaction between ROI lateralization and ROI (F (9) = 110.433, p<0.001). Pairwise comparisons were calculated as a post-hoc analysis (Table 2). After applying Bonferroni correction, all regions appeared to be significantly different (p<0.001). Almost all regions had more volume in the right hemisphere (contralateral to the language-dominant hemisphere). Only 2 structures were found to be bigger in the left hemisphere: the pars triangularis and middle temporal gyrus.

Both groups of patients and the healthy controls showed similar relations of volumetric differences between the left and right ROIs, as can be seen in Figure 3. The pars triangularis exhibited the same pattern for all patients and healthy controls: more volume in the left hemisphere. The middle temporal gyrus was bigger in the left hemisphere for the right tumor group and healthy controls but not for the left tumor group. The remaining regions were all bigger in the right hemisphere for all participants, except for the pars orbitalis which was bigger in the left hemisphere for the right tumor group. Healthy participants show a complete asymmetrical lateralization pattern whereas brain tumor patients do not present asymmetries for some of the regions. For the left tumor group, 4 out of the 10 selected regions were different when compared to the control group: the pars orbitalis, middle frontal gyrus, superior temporal pole, and middle temporal gyrus. In the case of the right tumor group, 7 regions were different from those of healthy controls: the pars orbitalis (bigger in the left instead of the right), pars opercularis, middle frontal gyrus, middle temporal pole, superior temporal pole, superior temporal gyrus and supramarginal gyrus.

## Discussion

In the present VBM study, we investigated the structural flexibility of the language network in patients with brain tumors affecting the language-dominant hemisphere. To identify structural changes dependent on tumor laterality we analyzed 2 groups of patients: patients with left tumors and patients with right tumors. Furthermore, a group of healthy controls was included to provide the standard lateralization pattern of the language network to enable the interpretation of the potential compensatory effects triggered by the growth of a brain lesion.

Three main findings can be highlighted, which we summarize here and discuss in more detail below. First, all of the patients, regardless of tumor laterality, showed a global change in the left language-dominant network and its contralateral counterpart compared to the control group. Second, contrary to what had been expected, a brain tumor induces compensatory neuroplastic mechanisms in both patient groups (left and right tumors), not just in patients with tumors in the left language-dominant hemisphere. Third, both tumor groups displayed different patterns of regional structural lateralization. Overall, these findings suggest that the growth of a brain tumor induces compensatory neuroplastic mechanisms in language regions that are not limited to patients with tumors in the left language-dominant hemisphere, with some specificities depending on tumor lateralization.

Our first main finding was GM dissimilarities between patients and healthy controls most likely induced by the presence of the tumor. This result is consistent with the previous findings focused on structural changes of the contralesional language counterpart (Almairac, Duffau, and Herbet 2018; Hu et al. 2020; Yuan et al. 2020). Furthermore, our results demonstrate that structural reshaping is not circumscribed to areas contralesional to the left language network, but instead, it spreads across both hemispheres. The entire network seems to change to cope with damage and maintain its functions. In spite of the limitations of the VBM method to explain the physiology underlying the neuroplastic mechanisms for recovery, a glioma can be considered to cause metabolic stress. Therefore, it is plausible to hypothesize that is removal might induce neuronal plasticity. As has been suggested before, this process is probably accompanied by secondary synaptogenesis and dendritic sprouting that might affect all regions within a network (Majewska, Newton, and Sur 2006). Likewise, cerebrovascular reactivity (CVR) could also be affected. In fact, impaired CVR has been found within the lesion and in the whole brain for patients with diffuse glioma (Fierstra et al. 2018).

Second, although structural changes in the cortex were expected only in the left tumor group, both tumor groups showed a similar structural pattern in which left and right ROIs were affected regardless of tumor location. Therefore, our results concur with recent studies that propose a shift from the traditional view of language being strongly left-lateralized, with the right hemisphere merely supporting it (Vigneau et al. 2011). Instead, the right hemisphere has been demonstrated to be actively involved in language functions, including second language learning (Vingerhoets et al. 2003; Park, Badzakova-Trajkov, and Waldie 2012) and language recovery after a brain lesion (Hugues Duffau, Denvil, and Capelle 2002; H. Duffau et al. 2003; Hope et al. 2017). As neuroplasticity seems to affect the language network (including the right non-dominant hemisphere), additional attention should be given to the right hemisphere in the assessment of language from a clinical standpoint. This idea challenges the current clinical standards in which intraoperative brain mapping in patients with brain tumors in the right hemisphere only accounts for social, somatosensory and visuospatial processes (Duchaine and Yovel 2015; Molenberghs et al. 2012; Bonini 2017; Schurz et al. 2014). Ignoring the right hemisphere’s contributions to language increases the risk of language deficits after surgery, as reported in previous studies (Vilasboas, Herbet, and Duffau 2017). Based on our findings, we predict a better recovery prognosis for those patients whose structural lateralization patterns within each region resemble those of the healthy controls. Ultimately, furthering our understanding of the global macrostructural reshaping of the language network in patients with brain tumors relative to healthy controls may help target future approaches to language therapy.

Our third main finding was that there were commonalities and differences in the structural relationships between homologous regions for both tumor groups in contrast to the healthy controls. With respect to the lateralization pattern in healthy controls, the right structures showed greater volume than the left counterparts, with the exception of the pars triangularis within the IFG and the middle temporal gyrus. The leftward asymmetry of these two critical language hubs has been well documented in postmortem and *in vivo* specimens (Foundas et al. 1996; Geschwind and Levitsky 1968; Wada, Clarke, and Hamm 1975). Nonetheless, the rightward or leftward asymmetry pattern of each of the regions is still under debate and actively being studied (Kong et al. 2018; Sha et al. 2021). Asymmetries are a core element of the typical organization of the brain. Cortical asymmetries have been linked to neuropsychiatric and cognitive disorders (Karolis, Corbetta, and Thiebaut de Schotten 2019; Okada et al. 2016). In addition, to evaluate the relationship between the structural asymmetry of the middle temporal gyrus and the functional language laterality, Reynolds et al. (2019) investigated the structural and functional development of the language asymmetry in 117 healthy children across early childhood. They demonstrated that the macrostructural asymmetry of the arcuate fasciculus – the WM tract connecting the IFG and the middle temporal gyrus – is pronounced from the age of 2 years and increases even more over time (Reynolds et al. 2019). Likewise, language models propose the IFG to be left lateralized whereas other regions are more bilateral (Hickok and Poeppel 2007). In our study, healthy participants showed asymmetries for all the regions whereas patients did not. However, language hubs considered crucial for maintaining the network topology, such as the pars triangularis within the IFG and the angular gyrus (Ius et al. 2011), exhibited the same pattern in all patients and healthy controls, demonstrating the capacity of the brain to change to accommodate language functions. This work demonstrates that, in addition to the previously documented functional reshaping in individuals with brain tumors when language hubs are damaged (Ille et al. 2019; Krieg et al. 2013; Quiñones et al. 2021; Połczyńska et al. 2021), structural changes also occur in both patients with tumors in the left (language-dominant) hemisphere and in patients with tumors in the right hemisphere. Further, we demonstrated these changes with GM volume indexes as GM has been stated to have great plastic potential (Sarubbo et al. 2015; Ius et al. 2011). These changes are likely representative of the compensatory mechanisms. On a similar note, structural changes should also be studied in different populations, such as stroke patients, as previous functional evidence has shown that the location of the lesion determines differential involvement of ipsilesional or contralesional areas (Stockert et al. 2020).

### Limitations and future directions

*T*hree limitations of the present work should be addressed. First, our sample size was limited by our access to data from this specific population, which constrains the amount of variability among patients. However, to mitigate the potential effects of confounding variables, we included sex, age, tumor type, and tumor size as nuisance covariates. Yet, they may have independent effects which cannot be disambiguated from those related to the sample size. To solve this, larger cohorts are needed to replicate the findings, increase statistical power and sensitivity, detect group differences and investigate possible relations with inter-individual variables, perhaps via large-scale multicenter collaborations. Larger samples would also aid in the development of guidelines for the presurgical assessment of patients with brain tumors affecting not only the left-language dominant hemisphere but also its right counterpart.

The second limitation is the cross-sectional nature of the current study, which does not allow examining how neuroplastic mechanisms unfold over time in individuals with tumors. Although cross-sectional studies provide useful contributions and clarifications (Pantelis et al. 2005) and led us to detect global changes most likely caused by the tumor growth over time, longitudinal studies are also needed. Such studies would result in direct and reliable evidence on how this neuroplastic mechanism occurs in each individual. This would help delineate neuroplasticity mechanisms for recovery and facilitate the identification of neuroimaging predictors for post-operative prognosis. The use of connectivity measures in future studies is fundamental to disentangle 1) whether only regions that are interconnected structurally and/or functionally follow the same neuroplastic patterns overtime and 2) if those patterns respond to fluctuations in the entropy of the system as suggested by the global difference encountered in our study.

The third limitation was that patients and healthy participants were collected in different centers, which prevented us from making a formal comparison. In the future, it would be desirable to test both patients and healthy participants in the same center to avoid any potential confounds.

## Conclusions

To our knowledge, this research is the first to show that a brain tumor affecting the left language network or its right homologue induces global structural reshaping, highlighting the brain’s plasticity. Our work emphasizes the need to extend the scope of presurgical and intraoperative brain mapping in patients with tumors, since the impact of a brain lesion appears to be more global. Intraoperative mapping should be designed to respect the anatomical substrate that is already going through neuroplastic processes to promote recovery and ultimately minimize long-term deficits. Future, multi-center longitudinal studies regarding the impact of a brain tumor on the neuroanatomy of language are needed to broaden our understanding of the processes of the structural and functional compensation in individuals with brain tumors.

## Data Availability Statements

The data are not publicly available due to the data-sharing policies of the different institutions involved concerning vulnerable clinical information. Codes for performing VBM are shared in github: https://github.com/lmansoo

## Acknowledgments

This research was supported by: the Ikerbasque Foundation; the Basque Government through the BERC 2022 2025 program; the Spanish State Research Agency through BCBL Severo Ochoa excellence accreditation CEX2020-001010-S; the Fundación Científica AECC (FCAECC) through the project PROYE20005CARR; Spanish Ministry of Economy and Competitiveness through the Plan Nacional RTI2018 093547 B I00 (*LANGCONN*) awarded to MC and IQ; the Spanish Ministry of Science, Innovation and Universities; the Fondo Social Europeo through the predoctoral grant PRE2019-091492 awarded to LM; and the NIH (NIDCD) Grant K01DC016904 (*Comprehensive pre-surgical identification of the critical language network in tumor patients)* awarded *to MP*. In addition, the authors would like to thank all the patients who agreed to take part in this study. Last but not least, we would like to thank Ms. Samiha Molla, University of California, Los Angeles, for her help with editing this work.

## Notes

### Competing Interest Statement

The authors have declared no competing interest.

